# Sequence Homology and Tissue Tropism Determine Superinfection Exclusion of Zika virus in *Aedes aegypti* Mediated by an insect-specific Binjari-Zika Virus Chimera

**DOI:** 10.64898/2026.08.04.742677

**Authors:** Wessel Willemsen, Alyssa J. Peterson, Marleen Henkens, Hans M. Smid, Hayden J. Rohlf, Tessa M. Visser, Constantianus J.M. Koenraadt, Roy A. Hall, Monique van Oers, Jody Hobson-Peters, Gorben P. Pijlman, Jessica J. Harrison, Leon E. Hugo, Jelke J. Fros

**Author notes:** Wessel Willemsen and Alyssa J. Peterson contributed equally to this work. Author order was determined by mutual agreement.

## Abstract

Arboviruses such as dengue, Zika, and chikungunya viruses cause widespread disease and continue to expand their geographical range due to climate change and vector spread. Insect-specific flaviviruses (ISFs) are promising biocontrol candidates of arboviruses, due to recent studies showing that prior infection with an ISF can reduce arbovirus replication in mosquitoes through superinfection exclusion (SIE). However, the route of infection, tissue tropism, pathogenesis and the mechanisms underlying SIE of ISFs in mosquitoes remain unclear. RNA interference (RNAi) is a potent antiviral response in insects, therefore it is expected that sequence homology between the ISF and the arbovirus will strengthen SIE. Here, we used ISF Binjari virus and a chimera containing the Zika virus structural proteins prME (BinJ-ZIKV) as a model system. Intrathoracic injection of BinJ-ZIKV in *Aedes aegypti* led to rapid systemic infection that excluded the midgut, subsequently blocking ZIKV dissemination from the midgut. SIE was strongest in tissues where primary-virus replication was highest. This spatial component of SIE was stronger when there was sequence homology between the ISF and arbovirus and displayed a strong 21nt siRNA response, suggesting RNAi contributed to the observed SIE. Upon oral inoculation, BinJ-ZIKV replicated efficiently in mosquitoes, was detected across multiple tissues, and saliva. BinJ-ZIKV also had higher infection establishment than BinJV at lower oral titres. SIE was observed for BinJ-ZIKV infection after oral exposure interfered with subsequent ZIKV midgut infection. Together, these findings support engineered ISF-chimeras as valuable experimental tools to dissect viral determinants of SIE and to optimize mosquito-based arbovirus interference strategies.

**Importance:** Annually, over 400 million people are infected with mosquito-transmitted viruses. Insect-specific flaviviruses (ISFs) can interfere with the transmission of clinically important viruses through a phenomenon termed superinfection exclusion (SIE). However, the mechanisms of SIE remain poorly understood. Using a Binjari virus chimera expressing Zika virus (ZIKV) structural proteins, we show that SIE is highly tissue-specific, with exclusion of ZIKV only occurring at sites where the chimera actively replicates and induces the mosquito antiviral RNA interference pathway. We further demonstrate that incorporation of Zika virus prM and E proteins into the ISF backbone enhances infection of the mosquito midgut following oral exposure, which enables direct inhibition of ZIKV infection after a subsequent infectious blood meal. Together, these findings define a replication-dependent, tissue-specific mechanism of ISF-mediated protection and provide a framework for reducing mosquito-borne virus transmission through SIE.

## Introduction

Arthropod-borne viruses (arboviruses) are transmitted by blood-feeding arthropod vectors such as mosquitoes, ticks, sandflies and midges. These viruses are significant drivers of the global disease burden, with over 390 million annual dengue virus (DENV) infections alone (1, 2). In recent years, arboviruses have expanded their geographical range due to urbanization, human mobility and climate change, exemplified by increased activity of West Nile virus in central and northern Europe where this virus is transmitted by indigenous *Culex spp.* Mosquitoes (3–9) and the global spread of the invasive mosquito species *Aedes aegypti* and *Ae. albopictus*, which are key vectors for DENV, Zika (ZIKV), chikungunya, and yellow fever (YFV) viruses,(10, 11).

The genus *Orthoflavivirus* (family: *Flaviviridae*) constitute the largest group of arboviruses that contribute to the global disease burden (12, 13). Flaviviruses are single- stranded, positive-sense RNA viruses. Their 11 kb genome codes for a single polyprotein that is cleaved into three structural proteins; capsid (C), pre-membrane/membrane (prM), and envelope (E); and seven non-structural proteins (14). In addition to the arboviruses, numerous flaviviruses that are restricted to insects have been described. These are collectively known as insect-specific flaviviruses (ISFs) (15–18). ISFs can modulate a mosquito vector’s susceptibility for medically important arboviruses through superinfection exclusion (SIE), wherein a primary viral infection inhibits the establishment of a secondary virus (16, 19–22). This interaction has prompted investigation of ISFs as potential biocontrol agents for arboviruses (20).

Unlike arboviruses, ISFs are maintained exclusively within mosquito populations, primarily through vertical transmission from infected females to their progeny (15, 16, 22, 23). This mosquito-restricted transmission provides a comparative framework for investigating viral and host determinants that govern infection, transmission, and propagation across vector- borne and vector-restricted systems. Although SIE has been reported for ISFs in cell culture systems(19, 24–26) and *in vivo* within mosquito hosts(24, 27–29), the underlying viral and vectorial determinants of SIE remain unknown. In particular, both the tissue tropism of the viruses involved and the extent to which these effects can be moderated through the activation of vector immunity and chimeric virus design are not well defined.

Chimeric ISFs are useful tools to dissect these mechanisms by enabling targeted replacement of viral structural components while retaining a mosquito-restricted replication phenotype through the non-structural proteins. An excellent candidate for this chimeric approach is the Binjari virus (BinJV), a lineage II ISF originally isolated from *Aedes normanensis* mosquitoes in Australia(30). BinJV has been proven to be extremely tolerable for the exchange of its structural proteins with those of pathogenic flaviviruses. Crucially, even when expressing foreign envelope proteins, BinJV chimeras retain a strict insect-specific phenotype and cannot infect vertebrate hosts (31). Here, we provide novel insights in how the ISF BinJV and a BinJ-ZIKV chimera replicate, disseminate and activate immunity in vector mosquitoes and we assess the tissue tropism and immune activation required for effective SIE by subsequent ZIKV challenge.

## Materials and Methods

### Cell culture

C6/36 cells (*Aedes albopictus*) were cultured at 27 °C in Leibovitz L-15 medium (Gibco, Carlsbad, CA, USA) containing 10% heat inactivated fetal bovine serum (FBS; Gibco), 2% tryptose phosphate broth (Gibco) and 1% nonessential amino acids (Gibco). Cells were mechanically detached and passaged twice a week. Vero E6 cells (*Chlorocebus aethiops*) were cultured in Dulbecco’s Modified Eagle Medium (DMEM; Gibco) supplemented with 10 % FBS and 100 units/ml Penicillin and Streptomycin (P/S; Gibco). Vero E6 cells were cultured at 37 °C and 5% CO_2_ and passaged twice a week after detaching the monolayer with Trypsin-EDTA (Gibco).

### Viruses

In this study, we employed a previously developed BinJV-based platform(31) in which the prME proteins (amino acid Ala123 to Ala794) of BinJV (GenBank ID: <u>MG587038</u>) were replaced with those of ZIKV_Natal(32)_ (GenBank ID: <u>KU527068</u>), generating the BinJ-ZIKV chimera (31). Superinfections were performed with ZIKV (GenBank ID: <u>KU937936.1</u>). Viral passage histories were described previously (26).

### Mosquitoes

*Ae. aegypti* mosquitoes (Rockefeller strain obtained from Bayer AG) were maintained as described previously (33). The Rockefellar strain was used in all experiments except whole-mosquito IFAs, which were performed using *Ae. aegypti* females derived from a colony originally established by QIMR Berghofer from mosquitoes collected in Innisfail, Queensland, Australia and maintained under identical conditions.

### Intrathoracic injection

*Ae. aegypti* mosquitoes were anaesthetized with 100% CO_2,_ and were transferred to a CO_2_ pad. Female mosquitoes were selected and injected with 69nl of BinJV, BinJ-ZIKV, or phosphate-buffered saline (PBS). Viral titres were diluted with Leibovitz cell culture media and administered at a concentration of 1 x 10^8^ 50% tissue culture infectious dose (TCID_50_)/ml resulting in 6.9 × 10^3^ TCID₅₀/mosquito.

### Mosquito survival assay

*Ae. aegypti* mosquitoes were Intrathoracically injected with BinJV, the BinJ-ZIKV chimera or PBS. An additional control group received CO₂ treatment only. Mosquitoes were provided with 6 % *ad libitum* glucose solution and survival was monitored daily.

### Oral infections

24 hours prior to blood feeding the glucose solution was replaced with water. Mosquitoes were subsequently offered infectious blood meals containing 2 x 10^6^ or 2 x 10^7^ TCID_50_/ml of BinJV, BinJ-ZIKV or ZIKV through Parafilm using the Hemotek PS5 feeder. Mosquitoes were allowed to feed for one hour in light conditions at 60% RH and 26 °C. Following blood feeding, mosquitoes were anaesthetised with 100% CO_2_ and fully engorged females were selected. During a 7-day incubation period following the initial infection, mosquitoes were allowed to lay eggs in the wet cotton used for sugar feeding on top of the buckets. Sequential blood meals were offered using the Hemotek PS5 feeder. Four days after the second blood meal, whole mosquitoes were collected and stored at -80 °C.

### Salivation assay

Mosquito saliva was collected as described previously with a few minor adaptations (33). Mosquitoes were anaesthetised with 100 % CO₂ and immobilised by removing their legs and wings with forceps, which were added to 1.5 ml Safe-Seal micro tubes (Sarstedt, Nümbrecht, Germany) containing ∼15 0.5 mm zirconium beads (Next Advance, Averill Park, NY, United States). The proboscis of each mosquito was inserted into a 200 μl pipet tip (Greiner Bio-One) containing five μl of a 1:1 solution of 50% glucose solution and FBS for 45 min. After salivation, the mosquito bodies were added to 1.5 ml Safe-Seal micro tubes containing zirconium beads. Saliva samples were added to 1.5 ml microtubes (Sarstedt) containing fungizone (2.5 μg/ml; Invitrogen), P/S (Penicillin 100 units/ml, Streptomycin 100 μg/ml) and gentamycin (50 μg/ml; Life Technologies) in either either 55 μl DMEM or Leibovitz cell culture media for the detection of ZIKV on Vero cells and BinJV or BinJ-ZIKV on C6/36 cells, respectively. Samples were stored at -80 °C (33). The infection prevalence in mosquito bodies and saliva were determined as previously described (34).

### Virus titration

Endpoint dilution assays (EPDA) were utilized to establish the tissue culture infectious dose 50% (TCID_50_). Dilutions (10-fold) of ZIKV or BinJV and Binj-ZIKV were prepared in DMEM or Leibovitz cell culture media, respectively. Cell culture media were supplemented with fungizone, gentamycin and P/S. Vero E6 cells from confluent tissue cultures were detached and dissolved in a total volume equal to 1 ml/1.67 cm^2^. C6/36 cells were detached from a confluent monolayer and 4 x diluted in Leibovitz medium. Cell suspensions were added to the virus dilutions in a 1:1 ratio, mixed and plated in 6-fold onto microtiter plates (Nunc, Sigma-Aldrich, Zwijndrecht, The Netherlands) (33). ZIKV titers were determined by CPE at 4 dpi and BinJV and BinJ-ZIKV titers at 7 dpi.

### Midgut dissection and whole-mount immunofluorescence staining

Whole-mount immunofluorescence staining of midguts was adapted from a protocol originally developed for *Pieris brassicae* caterpillars (35). Mosquito midguts were dissected in 10 µl of 1× PBS. Blood was removed by dipping individual midguts in 50 µl Milli-Q water. Pools of five midguts were placed in 1.5 ml microcentrifuge tubes and washed three times in 1× PBS. Viruses were inactivated in 200 µl a 4% formaldehyde per tube for 24 h at 4 °C. Midguts were then washed three times in 1× PBS and permeabilized by dehydration through a graded methanol series (70%, 80%, 90%, 96%, and 100 % methanol; 1 min each), followed by rehydration (96%, 90%, 80%, and 70% methanol) and transfer into PBS-T (1 × PBS containing 0.2 % Triton X-100). Incubation and wash steps were performed under mild agitation on a slowly rotating platform. Samples were washed three times for 20 min in PBS-T and blocked for 1 h at room temperature in PBS-T-B (1 × PBS containing 0.2% Triton X-100 and 1 % bovine serum albumin, BSA, lyophilized powder, crystallized, ≥98.0%; Sigma-Aldrich, Zwijndrecht, The Netherlands; cat. no. 05470). Midguts were incubated with murine-4G4 (36) (1:10 in PBS-T-B) for 24 h at 4 °C, followed by six, 20 minute washes in PBS-T. Next, tissues were incubated overnight at 4 °C with goat-antimouse Alexa Fluor 488 (Molecular Probes, Invitrogen, Eugene, OR; Cat. no. A11008, RRID:AB_143165) (1:300) together with TO-PRO- 3 iodide (Thermo Fisher Scientific, USA)(1:2000), in PBS-T-B. Samples were then washed three times in PBS-T and three times in PBS. Midguts were dehydrated through an ethanol series (30%, 50%, 70%, 80%, 90%, 96%, 100%, 100%, and 100%), cleared in 1:1, ethanol : xylene followed by pure xylene, and mounted in DePeX (Sigma-Aldrich, Darmstadt, Germany). Confocal microscopy was performed with a Leica Stellaris 5 confocal microscope (Leica, Germany), using a 40× lens, APO objective NA1.4 and a spectrally flexible white light laser.

### IFA of whole mosquito

BinJV NS1 was detected in mid-sagittal sections of mosquitoes by IFA using established protocols (37, 38). Briefly, legs and wings were removed and mosquito bodies were fixed in 4% PFA/0.5% Triton X overnight before they were transferred to 70% ethanol. Mosquitoes were dehydrated through ascending graded alcohol and imbedded in paraffin. Mid-sagittal paraffin sections (4 µM) were cut and fixed to positively charged slides, dried overnight at 37 °C. Paraffin-embedded mosquito sections were dewaxed and rehydrated through xylene and a descending graded ethanol series to water. Samples were incubated with blocking buffer (Biocare Medical Background Sniper, 2% BSA) for 15 minutes at room temperature. Next, sections were incubated with undiluted primary antibody (clone 4G4(39)) overnight at 4 °C in a humidified chamber. Sections were then incubated with Alexa Fluor 488–conjugated anti-mouse secondary antibody diluted 1:300 in Tris-buffered saline for 2 hours at room temperature in the dark followed by 4′,6-diamidino-2-phenylindole (DAPI) for 3 minutes. Sections were mounted using Dako fluorescent mounting medium. Fluorescence images where acquired using an Axioscan 7 slide scanner (Zeiss, Germany) using a 20x Plan-Apochromat objective lens (air, 0.8 NA) and an Axiocam 712 mono camera. Images were processed for presentation by adjusting brightness and contrast using ZEN 3.12 software (Zeiss, Germany) (37, 38).

### RNA Extraction

Individual mosquitoes that were stored at -80 °C in 1.5 ml Safe-Seal micro tubes containing zirconium beads were homogenized using a Bullet Blender storm (Next advance) for 1 minute, followed by the addition of 100 µL of Leibovitz cell culture media and blending for 2 additional minutes. 750 µL of TRIzol reagent (Invitrogen) was then added, and the samples were vortexed for 15 seconds, incubated at room temperature for 3 minutes, and stored at -20 °C overnight. After thawing and subsequent incubation at room temperature for 3 minutes, RNA was isolated following the manufacturer’s protocol.

### Small RNA sequencing

Small RNA libraries were generated from ∼1 μg total RNA from carcasses or midguts on a DNBSEQ sequencing platform (BGI Group, Shenzhen, Guangdong, China). Trimmed single-end FASTQ reads were generated with an in-house filtering protocol of BGI. Analysis of sRNA-seq was performed as described earlier. Briefly, reads were mapped to the viral genome of BinJ-ZIKV with Bowtie2 version 2.5.4 allowing 1 mismatch with a seed length of 28. Size distributions of small RNAs and genome distributions of siRNAs and piRNAs were produced with in house R scripts (26).

### qRT-PCR

First-strand cDNA synthesis was performed using 1 μl of total RNA with SuperScript III (Invitrogen) and random hexamers (Roche), according to the manufacturer’s instructions. qPCR was performed with 5 µL PowerTrack SYBR Green qPCR MasterMix (Invitrogen), 3,5 µL RNase-free water, 1 µL of cDNA and 0,5 µL 10 µM forward and reverse primers of Oligonucleotide combinations 5’- TCCAGAGCATCCCTACAAGA-3’, 5’-GATCTCCTTGACAACGCCATTA-3’ (BinJV), 5’- TCAGGAGGTGGTGTTGAAGG-3’, 5’- GTGCCCTTTCTCCATTTGGT-3’ (ZIKV) and 5’-ATGGTTTTCGGATCAAAGGT-3’, 5’-CGATAGCCTTCTTGCTGTTG-3’ (RPS7). Each sample was run in two technical replicates. A sample was scored positive if both technical replicates had Ct values <35, with a between-replicate difference (ΔCt) of <0.5. Relative viral load was calculated using the ΔCt method with *RPS7* as the reference gene and expressed as 2^(-ΔCt).

### Statistics

For mosquitoes challenged with ZIKV after intrathoracic injection, infection, dissemination, and transmission rates were analysed using binomial logistic regression with replicate as a fixed blocking factor to account for between replicate variation. For each outcome, a full model (group + replicate) was compared to a replicate-only model via likelihood ratio test (LRT) to assess the overall group effect. Pairwise contrasts against the PBS control are reported as odds ratios (OR) with 95% Wald confidence intervals. Where complete or near-complete separation was detected, indicated by fitted probabilities of 0 or 1, non-finite coefficients, or standard errors exceeding 1000, models were refit using Firth’s bias-reduced logistic regression via the brglm2 package (brglmFit, AS_mean bias correction). This was necessary for the saliva outcome in both experiments due to the low number of positive cases. All analyses were performed in R using the glm function from base R and the brglm2, emmeans, and broom packages. The full test statistics (Table S1A,B), including the raw data (Table S2), are provide in the supplementary files.

All other statistical comparisons were conducted using GraphPad Prism 11.0.0. Mosquito survival rates were evaluated using the log-rank (Mantel-Cox) test. For continuous data including normalized viral RNA levels and infectious titers, non-parametric analyses were employed as the data did not meet the assumptions of normality. Comparisons between two independent groups were performed using the Mann-Whitney U test, while comparisons across more than two groups were analyzed via the Kruskal-Wallis test followed by Dunn’s multiple comparisons post-hoc test. Differences in viral prevalence between two specific treatment groups were assessed using Fisher’s exact test. For all analyses, a p-value of less than 0.05 was considered statistically significant.

## Results

### Replication of BinJV and BinJ-ZIKV has no effect on *Aedes aegypti* survival

To investigate viral replication dynamics and the effect of intrathoracic injections and virus infection on mosquito fitness, *Ae. aegypti* mosquitoes were injected with either PBS, BinJV or BinJ-ZIKV. No significant differences in mosquito survival were observed between treatment groups and a not injected control group (Ctrl) (log-rank test, p = 0.12) (Fig. 1A). Infection rates were 100% at all time points post injection. Three days post injection, normalized viral RNA levels increased approximately 1000-fold compared to the day of injection and remained high over time, with BinJV reaching higher normalized viral RNA levels compared to BinJ-ZIKV (Fig. 1B).

**Figure 1:**
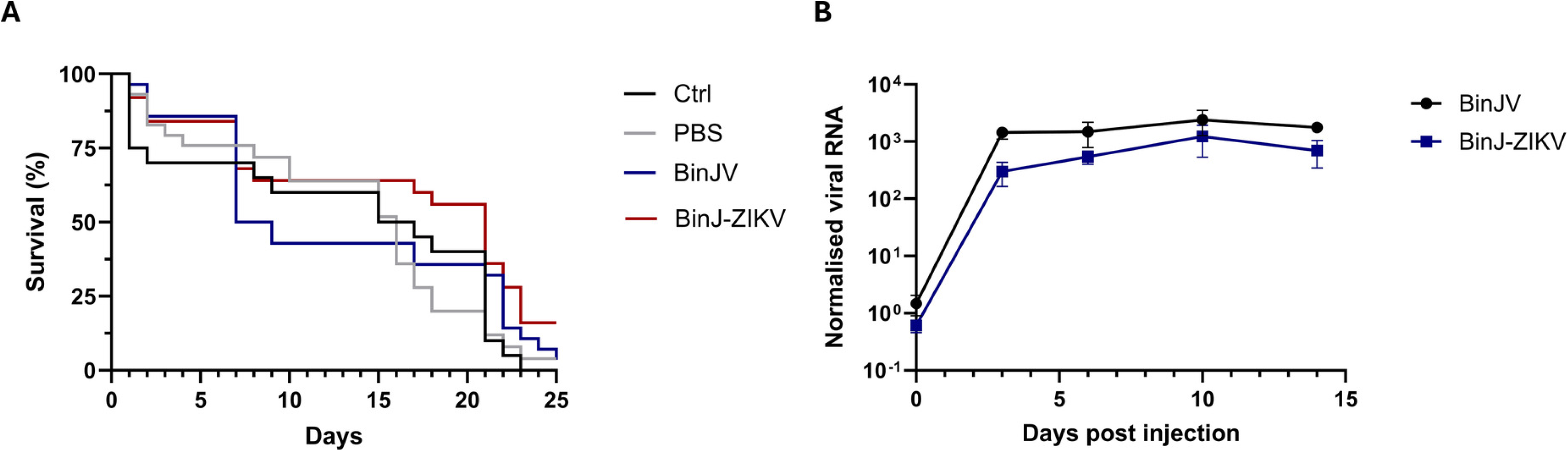
Infection dynamics of BinJV and BinJ-ZIKV in *Ae. Aegypti* after intrathoracic injection. A) Survival rates of non-injected control (Ctrl, n=20), PBS injected (n=26), BinJV injected (n=28), or BinJ-ZIKV (n=25) injected *Ae. aegypti* (log-rank test, p = 0.12). B) Viral RNA load from BinJV or BinJ-ZIKV normalised to the housekeeping gene *RPS7* from whole mosquitos following injection. Mean values are plotted with each point representing 5-9 mosquitoes with error bars showing standard error of the mean.

### Intrathoracic injection and BinJ-ZIKV infection specifically reduce *Aedes aegypti* susceptibility to ZIKV

Next, the inhibitory capacity of BinJV and BinJ-ZIKV in a mosquito vector was investigated. First, *Ae. aegypti* mosquitoes were injected with either PBS, BinJV or BinJ-ZIKV and offered an infectious blood meal containing 2×10^6^ TCID₅₀/ml ZIKV (Fig. S2). Generally, ZIKV infection rates were lower than expected given the appropriate virus titer in the blood meal based on previous ZIKV infections of *Ae. aegypti* in our lab (∼80-90%)(40). Although no ZIKV dissemination to the legs and wings was observed in the BinJ-ZIKV injected group compared to the PBS injected group (Fig. S2), these low overall rates across all groups prevented meaningful statistical comparisons (Table S1A,B). Importantly, because all ZIKV infection rates were low even in the PBS injected control group, we hypothesized that the injection itself might interfere with the ability of ZIKV to infect *Ae. aegypti*.

To test this hypothesis, we included a control group (Ctrl) which was anesthetized with CO_2_ for the same amount of time as the injected groups but received no injection, and offered mosquitoes an infectious blood meal containing a ten times higher dose of ZIKV at 2×10⁷ TCID₅₀/ml to increase infection prevalence for statistical power. For each mosquito, the legs and wings, saliva and the remainder of the carcass (still containing the midgut and hereafter called “body”), were isolated 10 days post blood meal and scored for the presence of ZIKV to determine the dissemination, transmission and overall infection rates (Fig. 2A).

**Figure 2.**
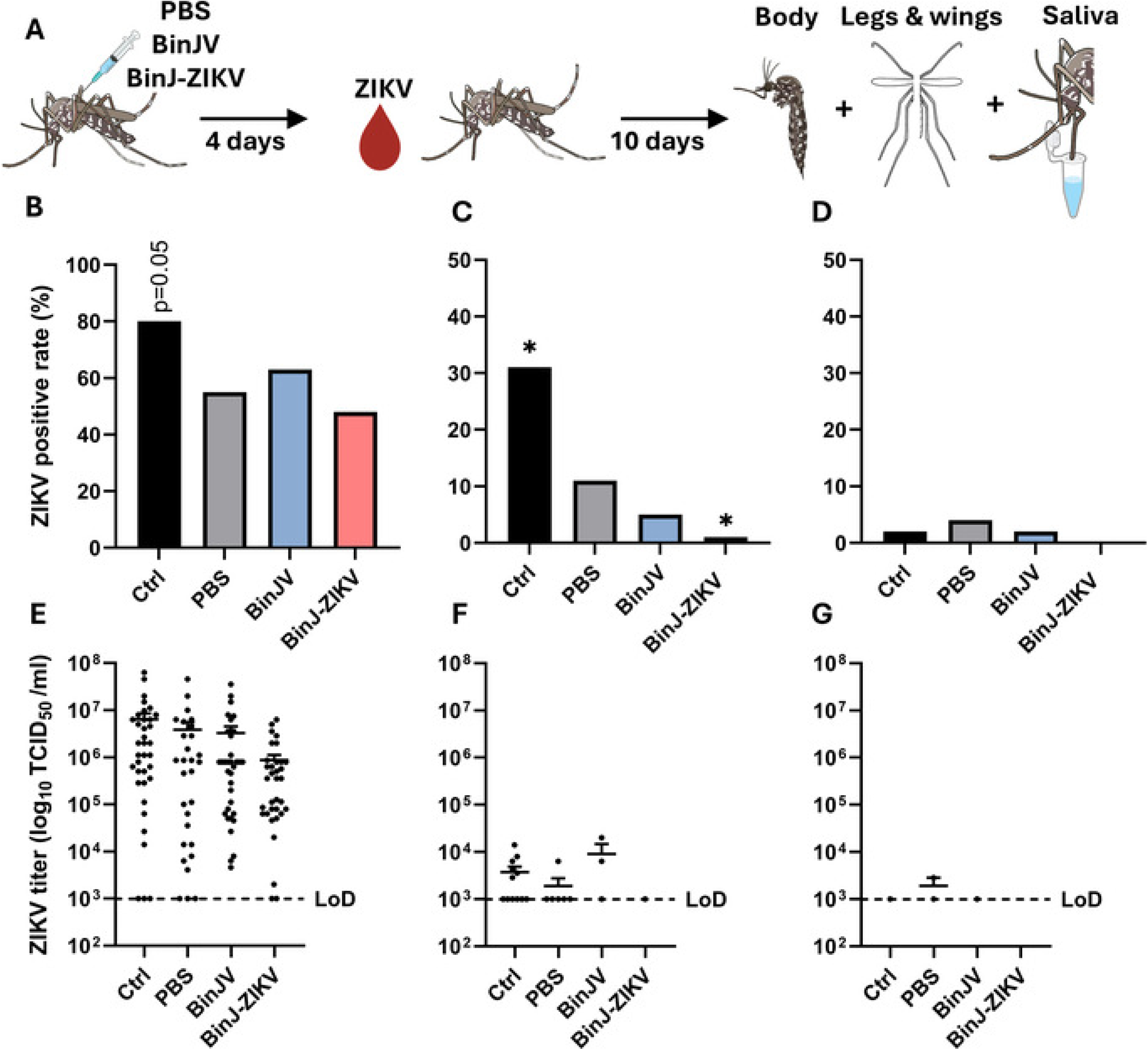
ZIKV infection and dissemination in mosquitoes injected with BinJV, BinJ-ZIKV. A) *Ae. aegypti* mosquitoes were intrathoracically injected with either PBS (n=55), BinJV (n=57) or BinJ-ZIKV(n=71) (6.9 × 10^3^ TCID₅₀/mosquito). A non-injected control group (Ctrl) was also included (n=45). Four days later, mosquitoes received a blood meal containing ZIKV. Ten days later, bodies, legs and wings, and saliva were collected and the ZIKV infection rate was determined. B-D) Percentage of mosquitoes positive for ZIKV in bodies (B), legs and wings (C), and saliva (D). E–G) Individual mosquito ZIKV titres in bodies (E), legs and wings (F) and saliva (G). Bars represent the average percentage of mosquitoes positive for ZIKV from three independent experiments. Asterisk indicates significant difference with the PBS injected group (GLM with replicate as blocking factor, p<0.05). Data points represent viral titers obtained from a single mosquito sample. Individual samples from three biological replicates are presented together with their respective mean and standard error of the mean. The dotted line indicates the limit of detection (LoD).

Interestingly, infection rates differed significantly across groups (GLM with replicate as blocking factor, overall group test, p = 0.0169). The Ctrl group had a higher infection rate compared to the PBS injected group (n = 55; GLM with replicate as blocking factor, OR = 2.62, p = 0.050), whereas neither BinJV (n = 57; OR = 1.16, p = 0.721) nor BinJ-ZIKV (n= 71; OR = 0.625, p = 0.234) differed significantly from the PBS injected group (n = 55) (Fig. 2B). Dissemination rates were also significantly different between treatments (GLM with replicate as blocking factor, overall group test, p = 1.33×10⁻⁵) (Fig. 2C). The non-injected control group had the highest dissemination rates (31%), significantly higher than the 10% observed in the PBS injected group (n = 55; GLM, OR = 3.13, p = 0.047). Conversely, the BinJ-ZIKV injected group had significantly lower dissemination rates at 1% (n = 71) (n = 55; GLM, OR = 0.091, p = 0.031).

Although very low potential ZIKV transmission rates were observed for all groups, without significant differences between groups (Firth/bias-reduced binomial model, overall group test, p = 0.553) (Fig. 2D), it is noteworthy that none of the mosquitoes injected with BinJ-ZIKV had ZIKV in the saliva (n = 71). Similarly, viral titers in the bodies were highest in the non-injected control group, lower in the PBS and BinJV injected groups, and lowest in the BinJ-ZIKV injected group (Fig. 2E). Compared to the PBS injected group, none of the groups differed significantly in body titers; however, BinJ-ZIKV did have significantly lower titers compared to the non-injected control group (Kruskal–Wallis with Dunn’s multiple comparisons, adjusted p = 0.024). Moreover, ZIKV titers in legs and wings were detectable in all groups except BinJ-ZIKV (Fig. 2F), whereas in saliva, viral titers were detectable only in the PBS-injected group (Fig. 2G).

These results indicate that the physical stress of the injection alone negatively impacts the vector competence of *Ae. aegypti* for ZIKV and that ZIKV dissemination was absent or strongly reduced in BinJ-ZIKV injected mosquitoes.

### Intrathoracic injection results in minimal BinJ-ZIKV replication and siRNA production in the *Aedes aegypti* midgut

Because BinJ-ZIKV reduced the ZIKV dissemination, but not the infection rate, we hypothesized that BinJ-ZIKV fails to productively infect midgut cells post intrathoracic injection limiting SIE of ZIKV. To determine how direct intrathoracic injection of BinJV or BinJ-ZIKV affects tissue distribution, specifically within the midgut, viral RNA in dissected midguts and remaining carcasses (without the midgut) was quantified at 4 dpi. Normalised viral BinJV and BinJ-ZIKV RNA levels were significantly higher in the carcass of injected mosquitoes compared to the midgut for both viruses (Fig. 3A) (Mann-Whitney U test, p<0.0001). Next, small RNA sequencing (sRNA-seq) was performed on the midgut and carcass at 4 dpi with BinJ-ZIKV (Fig. 3B) to assess activation of the antiviral RNAi response. A strong 21 nt siRNA peak was mounted against BinJ-ZIKV in the carcass, but not the midgut (Fig. 3B). The 21-nt siRNAs mapped to both sense and antisense strands of the entire BinJ-ZIKV genome, including the ZIKV prME genes (Fig. 3C).

**Figure 3:**
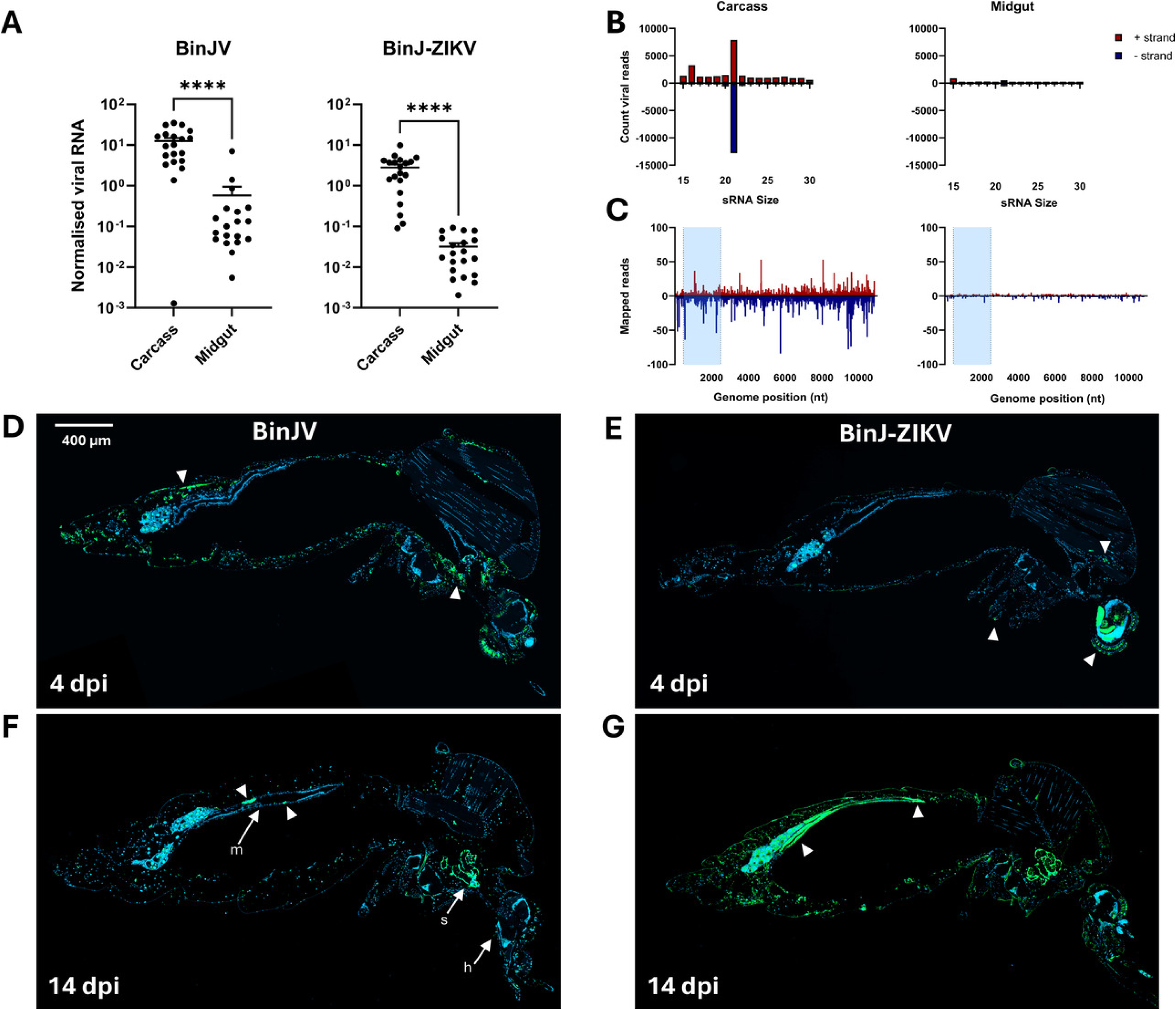
Tropism of BinJV and BinJ-ZIKV in *Ae. Aegypti* after intrathoracic injection. A) Viral RNA load of BinJV or BinJ-ZIKV normalised to the housekeeping gene *RPS7* in carcass vs. midgut at 4 dpi (n=20, Mann-Whitney U test, p<0.0001). B) Size distribution of 15-30 nt small RNAs that map to the genome of BinJ-ZIKV. isolated from either the carcass or midgut (n=10). C) genome distribution of 21 nt small RNAs that map to the genome of BinJ-ZIKV. Blue rectangle indicates the location of the ZIKV prME genes. D-G) Whole *Ae. aegypti* sections immunostained for flavivirus NS1 (green), and nuclei (blue). Panels show mosquitoes 4 days post-injection (dpi) with BinJV (D) or BinJ-ZIKV (E), and 14 dpi with BinJV (F) or BinJ-ZIKV (G). White arrowheads indicate green fluorescence representing BinJV or BinJ-ZIKV-related signals. White arrows in F illustrate the positions of head (h), salivary gland (s), and midgut (m).

Whole-mosquito IFA were performed on *Ae. aegypti* injected with BinJV and BinJ- ZIKV and stained for BinJV NS1 to visualise the viral tropism. At 4 days post-injection BinJV NS1 was primarily localised in the head and carcass, while remaining largely absent from the midgut for both BinJV (Fig. 3D) and BinJ-ZIKV (Fig. 3E). By 14 days post-injection, small focal points of NS1 were additionally observed in the midgut for the BinJV infected mosquito (Fig. 3F), whereas extensive infection spanning almost the entire midgut was evident in the BinJ-ZIKV injected mosquito. These results suggest that local siRNA responses are mounted quickly in infected tissues, while the mosquito midgut takes longer to establish robust replication and siRNA responses after intrathoracic injection.

### Oral inoculation of BinJ-ZIKV stimulates robust replication and RNAi activation in the midgut epithelium

Next, we tested whether both viruses could directly establish a potent infection in the midgut following an infectious blood meal. *Ae aegypti* were offered an infectious blood meal with either BinJV or BinJ-ZIKV at 2*10^6^ TCID_50_/ml. After 14 days, total RNA was extracted from whole mosquitoes and infections were examined by RT-qPCR (Fig. 4A). BinJ-ZIKV was significantly more effective at infecting *Ae aegypti* via infectious blood meal compared to BinJV with respectively 75% and 15% of mosquitoes testing positive (Fisher’s exact test, p < 0.0001) (Fig. 4B,C). After 14 days, the legs and wings were separated and saliva collected from each mosquito. The remaining body, legs and wings, and saliva samples were inoculated on permissive C6/36 cells to determine the infection, dissemination and transmission rates of BinJV and BinJ-ZIKV, respectively (Fig. 4A). Again, BinJ-ZIKV (n = 27) had significantly higher infection rate in the body (Fig. 4D,G) at 89% compared to BinJV (n = 15) at 13% (Fisher’s exact test, p < 0.0001). Consistent with this, a dissemination rate of 33% (Fig. 4E,H) was detected in BinJ-ZIKV–infected mosquitoes compared to 7% for BinJV, and virus was present in saliva in 22% of BinJ-ZIKV–infected mosquitoes compared to 7% for BinJV (Fig. 4F,I). This indicates that both viruses are able to infect, disseminate and localize to the saliva of *Ae. aegypti* after an infectious blood meal. However, BinJ-ZIKV consistently displayed higher percentages of infection compared to BinJV, suggesting BinJV was restricted at the initial stage of infecting the midgut.

**Figure 4.**
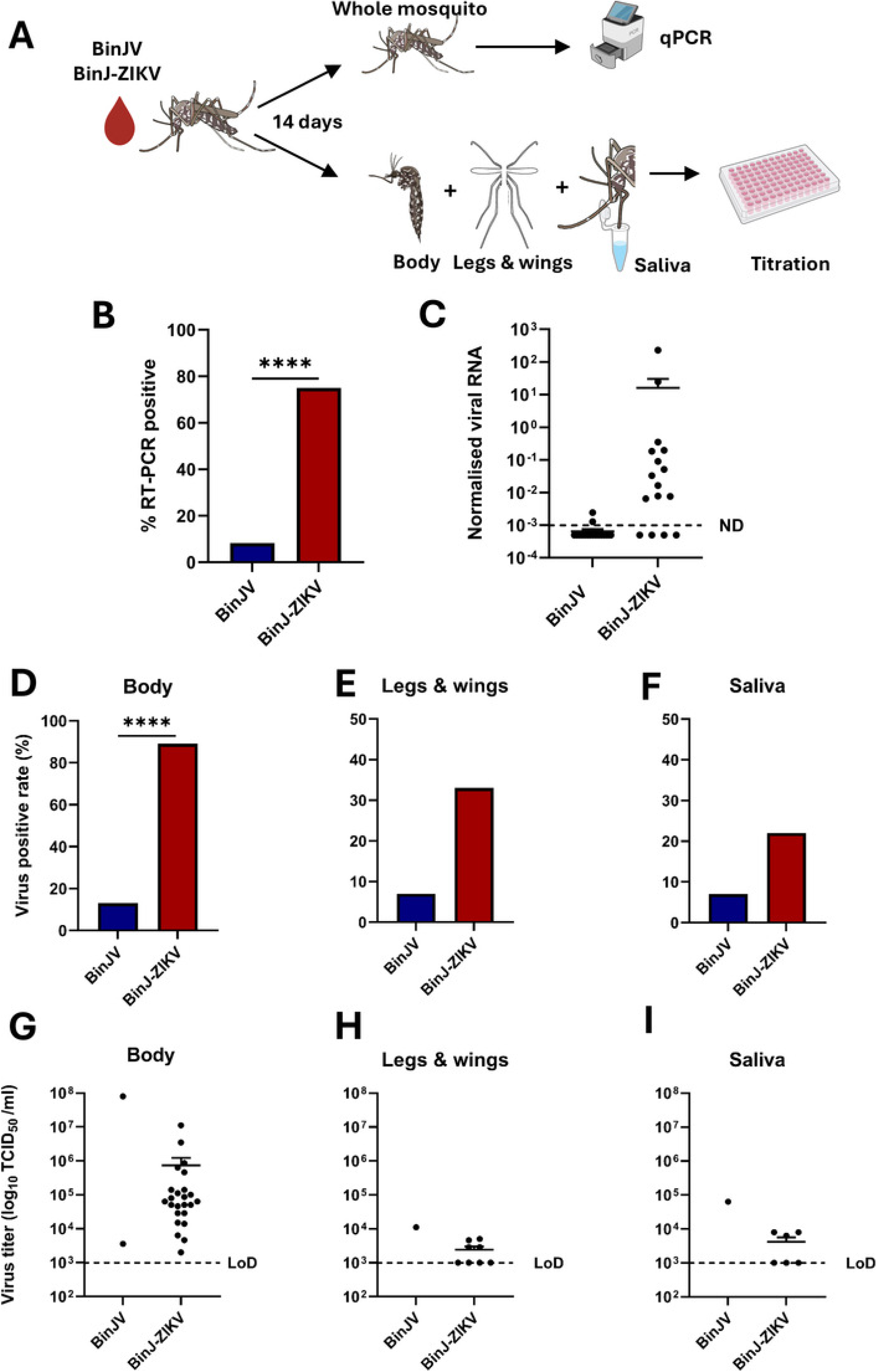
Infection, dissemination, and transmission efficiency of BinJV and BinJ-ZIKV in *Ae. aegypti* following oral infection. A) *Ae. aegypti* mosquitoes were offered an infectious blood meal containing BinJV or BinJ-ZIKV and analysed 14 days post-feeding. B) Percentage of mosquitoes positive for BinJV (n=23) or BinJ-ZIKV (n=16) by RT-PCR. C) Viral RNA load normalised to the housekeeping gene *RPS7* in individual whole mosquitoes. The dotted line indicates the detection limit below which virus was not detected (ND). D) Virus-positive rate in mosquito bodies (BinJV n=15, BinJ-ZIKV n = 27). E) Virus-positive rate in legs and wings (dissemination). F) Virus-positive rate in saliva (transmission). G-I) Infectious viral titres (log₁₀ TCID₅₀/mL) in mosquito bodies (G), legs and wings (H) and saliva (I). Dotted lines indicate the limit of detection (LoD) with error bars showing standard error of the mean. Asterisks indicate significant differences (Fisher’s exact test, p < 0.0001).

To further investigate the initial infection of the midgut, mosquitoes were sacrificed four days after receiving an infectious blood meal containing 2*10^6^ TCID_50_/ml of BinJV, BinJ- ZIKV or ZIKV as an oral infecting arbovirus control. Midguts were dissected and immunostained for flavivirus NS1. NS1 proteins of BinJV and ZIKV were readily detected in the midgut epithelium of *Ae. aegypti* four days post blood meals with BinJ-ZIKV and ZIKV, respectively (Fig. 5A). Blood meals containing BinJV did not yield any NS1 positive midguts (n=10). Mosquitoes exposed to BinJ-ZIKV, typically displayed a clear site of infection, consisting of a dense cluster of infected epithelial cells. ZIKV-infected mosquitoes displayed multiple infection foci distributed throughout the midgut (Figure 5B).

**Figure 5.**
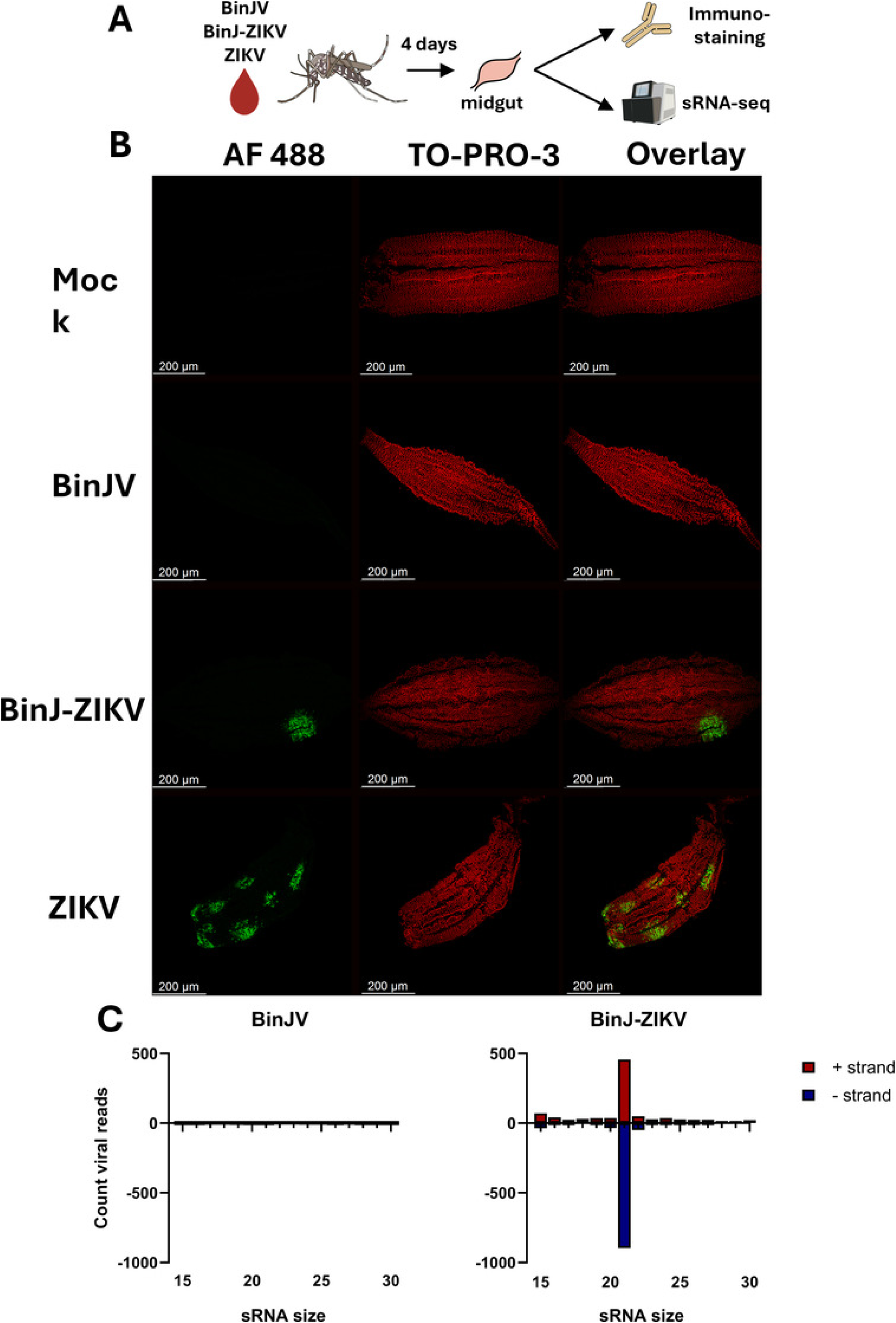
Characterisation of midgut infections following an infectious blood meal. A) *Ae. aegypti* mosquitoes were offered an infectious blood meal containing BinJV or BinJ-ZIKV after which their midguts were dissected and analysed 4 days post-feeding by either IFA or sRNA-seq. B) Representative confocal images of dissected midguts immunostained with anti-flavivirus NS1 monoclonal antibody 4G4 detected with Alexa Fluor 488 (green), counterstained with TO-PRO-3 to label nuclei (red), and merged images shown. Scale bars indicate 200 µm. C) small RNA sequencing size distribution of reads isolated from midguts (n=10) mapping to either BinJV or BinJ-ZIKV.

To further explore the antiviral siRNA response to BinJV or BinJ-ZIKV infection, sRNA-seq was performed on RNA isolated from 10 midguts exposed to either BinJV or BinJ- ZIKV. A strong antiviral RNA interference response was observed against BinJ-ZIKV, characterized by a prominent 21-nt peak mapping to both the sense and antisense strands of the entire chimeric viral genome (Fig. 5C). In contrast, hardly any siRNAs (7 reads) mapped to the BinJV genome in the mosquitoes that fed on BinJV containing blood.

Together, these results demonstrate that the structural prME proteins of ZIKV enable BinJ-ZIKV to enter and replicate within the midgut of *Ae. aegypti*. While BinJV itself is largely restricted from establishing infection via the oral route, incorporation of ZIKV structural genes appears to overcome this midgut barrier, facilitating the initial infection of the midgut and eliciting a strong siRNA response.

### Exclusion of superinfecting ZIKV within individual *Aedes aegypti*

To investigate whether BinJ-ZIKV infection of the midgut interferes with a secondary ZIKV infection, *Ae. aegypt*i were offered an infectious blood meal with either BinJV or BinJ-ZIKV at 2*10^7^ TCID_50_/ml. After 7 days a subsequent blood meal containing ZIKV at 2*10^6^ TCID_50_/ml was offered. Four days post ZIKV infection, whole mosquitoes were homogenized and processed to quantify infection virus and viral RNA (Fig 6A). Immunostaining of the mosquito midgut post primary oral infection with BinJ-ZIKV displayed that after 7 days a much larger portion of the midgut is infected compared to 4 days (Fig. 6B), potentially improving the exclusion of superinfecting ZIKV.

**Figure 6.**
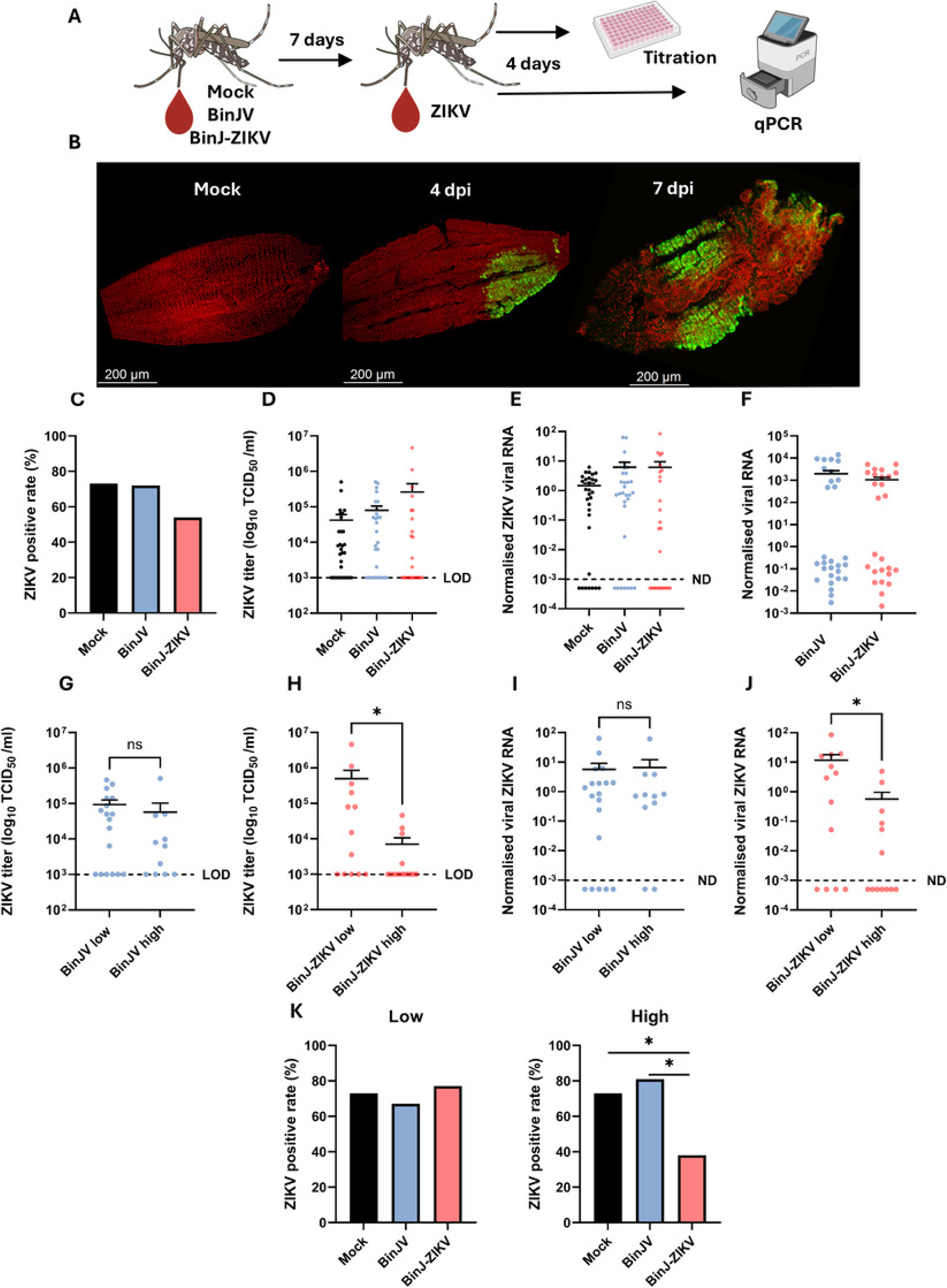
BinJ-ZIKV suppresses ZIKV infection in *Ae. Aegypti*. A) *Ae. aegypti* mosquitoes were given a blood meal containing BinJV or BinJ-ZIKV (2 × 10⁷ TCID₅₀/mL). Seven days later, mosquitoes received a second blood meal containing ZIKV (2 × 10⁶ TCID₅₀/mL). At 4 days post ZIKV infection, whole mosquitoes were collected for infectivity assays and qPCR analysis. B) Representative confocal images of dissected midguts. Infection is shown in green (NS1) and nuclei in red (TO-PRO-3). Images are shown for mock-treated mosquitoes and BinJ-ZIKV-treated mosquitoes at 4 and 7 dpi. C) Mosquitoes received a primary blood meal containing no virus (mock, n = 30), BinJV (n = 29), or BinJ-ZIKV (n = 26). Bars represent the percentage of mosquitoes positive for ZIKV after the second blood meal. D) Infectious ZIKV titres in mosquito bodies (log₁₀ TCID₅₀/mL). E,F) The ZIKV (E) and BinJV or BinJ-ZIKV (F) RNA load normalised to the housekeeping gene *RPS7.* G–H) Infectious ZIKV titres compared between mosquitoes containing low and high RNA levels of BinJV (G) and BinJ-ZIKV (H). I–J) ZIKV RNA load normalised to the housekeeping gene *RPS7* compared between mosquitoes containing low and high RNA levels of BinJV (I) and BinJ-ZIKV (J). Data points represent individual mosquitoes. Dashed line indicates the limit of detection (LoD) for viral titers or below which viral RNA was not detected (ND). Asterisks indicate significant differences (Mann–Whitney U test, p <= 0.05). *K)* Bars represent the percentage of ZIKV-positive mosquitoes when mosquitoes contained either low (left) or high (right) levels of BinJV or BinJ-ZIKV RNA. Asterisks indicate significant differences (Fisher’s exact test, p < 0.05).

After the ZIKV blood meal, ZIKV infection rates were very similar in the mock treated and BinJV treated groups at 73% and 72% respectively. The BinJ-ZIKV treated group had the lowest ZIKV infection rate at 54% (Fig. 6C). However, no differences were observed between the ZIKV titer or RNA levels between groups (Fig. 6D,E). Interestingly, when quantifying the levels of normalized BinJV and BinJ-ZIKV RNA, all mosquitoes had detectable RNA levels for both the BinJV and BinJ-ZIKV treated group (Fig. 6F). Notably, a bimodal distribution was observed within both the BinJV and BinJ-ZIKV treated mosquitoes, characterized by either high or low viral RNA levels. Approximately 38% and 50% contained high levels of RNA from BinJV and BinJ-ZIKV, respectively.

Subsequently, the corresponding ZIKV titers were split between the BinJV and BinJ- ZIKV low and high RNA level groups. For the BinJV infected mosquitoes no significant difference was observed between the groups (Fig. 6G). However, in the BinJ-ZIKV infected group significantly lower ZIKV titers (Mann-Whitney U test, p = 0.0387) were observed when mosquitoes carried high levels of BinJ-ZIKV RNA compared to low levels of RNA (Fig. 6H). Similarly, the presence of high levels of BinJV RNA did not change the levels of ZIKV RNA detected (Fig. 6I), whereas significantly lower ZIKV RNA levels were detected in mosquitoes carrying high levels of BinJ-ZIKV RNA (Mann-Whitney U test, p =0.0326) (Fig. 6J).

Finally, the ZIKV infection rate was divided between mosquitoes with low and high BinJV and BinJ-ZIKV RNA loads (Fig 6K). ZIKV prevalence was similar between all three groups in the low RNA level group at 73%, 67% and 77% for mock, BinJV and BinJ-ZIKV respectively. For the high RNA level group, the ZIKV prevalence was significantly lower if mosquitoes were infected with BinJ-ZIKV compared to BinJV (Fisher’s exact test, p = 0.0472) or the mock (Fisher’s exact test, p = 0.0427).

In summary, we revealed that a prior infection of *Ae. aegypti* with BinJV did not inhibit subsequent ZIKV infection. However, productive replication of the BinJ-ZIKV chimera in the midgut interferes with subsequent ZIKV infection and significantly lowered ZIKV titers, RNA levels, and infection prevalence, while mounting a strong siRNA response.

## Discussion

ISFs and chimeras between ISFs and arboviruses are increasingly recognised as promising tools for vector-borne disease biocontrol and have been used as platforms for recombinant vaccines and diagnostic antigen production against medically important orthoflaviviruses(41–45). Despite this growing translational interest, the *in vivo* dynamics of ISF infection and the mechanistic determinants of superinfection exclusion (SIE) in mosquitoes remain incompletely understood. To comprehensively evaluate SIE, we compared commonly used experimental routes of mosquito infection and tracked the subsequent viral kinetics. Specifically, we investigated the infection and dissemination dynamics within the vector, focusing closely on the midgut as a critical anatomical barrier. Utilizing these foundational insights, we present *in vivo* SIE studies that employ a dISF BinJV-based chimera expressing Zika virus (ZIKV) prM and E structural proteins in *Aedes aegypti*, a principal vector responsible for ZIKV transmission(46, 47).

Intrathoracic injection resulted in significantly higher viral RNA loads for both BinJV and BinJ-ZIKV in carcasses compared to midguts at four days post injection, and small RNA sequencing further confirmed active replication of the chimera, with abundant 21-nt virus-derived siRNAs. Whole-mosquito IFA staining further supported these findings, showing viral antigen localized predominantly in the mosquito body (carcass) and largely absent from the midgut. This is in contrast to intrathoracic injection of the ISF Palm Creek Virus (PCV) whereby the ISF was exclusively detected in the midgut(48), but consistent with recent research where microinjection of a BinJV reporter virus localized to the salivary gland and not the midgut 7 days post injection(49). Interestingly, 14 days post injection viral antigen was detected in the midgut for both BinJV and BinJ-ZIKV in this study, showing that incubation time and virus specific tissue tropism are critical factors to consider when evaluating *in vivo* SIE for biocontrol.

The injection-based SIE experiments revealed an important methodological limitation. Overall ZIKV superinfection rates were reduced across all injected groups, including PBS controls. Inclusion of an injection-free control demonstrated substantially higher ZIKV infection and dissemination in non-injected mosquitoes, indicating that the injection procedure itself likely induced an immune response that altered vector competence. Mechanical injury and systemic immune activation have previously been shown to influence arbovirus susceptibility in mosquitoes, particularly via RNAi and other innate immune pathways (50, 51). In a recent *in vitro* study(26), we noted that acute infection with BinJ-ZIKV induces severe cytopathology and cell death in two different siRNA deficient cell lines, which overestimates the true baseline of viral exclusion. These findings emphasise that both physical trauma from injection in live vectors and acute cellular stress *in vitro* represent experimental conditions that can confound SIE metrics. Thus, reduced arbovirus infection following intrathoracic manipulation cannot be interpreted solely as viral interference and highlight the need for caution when assessing SIE using injection-based models.

Despite the confounding effects of injection, BinJ-ZIKV infection was associated with the lowest levels of ZIKV dissemination. In particular, ZIKV detection in legs and wings was almost entirely absent in the BinJ-ZIKV group, suggesting that sequence homology in the prM and E region may promote tissue local interference once the chimera has established replication. This mirrors *in vitro* observations where RNAi-competent Aag2 cells pre-infected with BinJ-ZIKV demonstrated stronger exclusion of secondary ZIKV challenges than those pre-infected with BinJV(26). Furthermore, the observed tissue local pattern is consistent with prior work showing that injection of ZIKV defective viral genomes reduced viral dissemination without preventing initial midgut infection(52). Taken together this supports a model in which homologous interference is only present in tissues where the primary virus is actively replicating at the time of the superinfection.

Oral infection experiments revealed that BinJV was largely restricted by the midgut infection barrier in *Ae. aegypti* and suggested that the ZIKV prM and E structural proteins mediate efficient midgut cell entry. Similarly, CHAOV-derived chimeras expressing ZIKV prM/E, exhibited enhanced midgut cell infection and altered arbovirus transmission dynamics, suggesting the midgut infection barrier remains a primary determinant of successful virus establishment in mosquitoes (53, 54). When the infectious doses in blood were increased tenfold, both BinJV and BinJ-ZIKV infected mosquitoes following oral exposure. This indicates that structural-protein-mediated enhancement of midgut entry is most pronounced at lower infectious doses and becomes less discriminatory under high-dose conditions. A higher infectious dose may alter the concentrations of other inoculum-associated components, including viral proteins such as NS1 as culture supernatant was used as opposed to purified virus. NS1 has recently been shown to promote midgut infection establishment for DENV(55, 56). Whether BinJV NS1 is secreted efficiently from C6/36 cells or is released into the culture supernatant through cytopathic effect-mediated cell lysis, thereby contributing to infection of a mosquito midgut remains unknown and warrants further investigation.

Mosquitoes with high levels of BinJ-ZIKV replication exhibited reduced ZIKV titres, RNA loads, and infection prevalence, compared to those with lower replication levels. CHOAV/ZIKV infection similarly reduced ZIKV replication(53). Together, these findings indicate that effective SIE requires sufficient primary virus replication within the midgut and likely operates through local mechanisms. We observed clear production of virus specific siRNAs in those tissues that became refractory to ZIKV infection. However, we cannot exclude potential involvement of other factors, such as competition for susceptible cells, cellular resources, or sustained activation of additional antiviral pathways. Collectively, these results support a model in which incorporation of ZIKV structural proteins into a BinJV backbone helps to overcome the midgut infection barrier and to enable replication-dependent SIE of secondary ZIKV infection.

From a translational perspective, these findings provide both encouragement and caution. ISF-based chimeras demonstrate clear mechanistic potential for modulating arbovirus infection dynamics and theoretically provide a means to prevent arbovirus infections without medical interventions. However, achieving consistently high chimera replication across a mosquito population remains a significant challenge for potential biocontrol strategies. Ultimately, establishing ISFs at a population level depends on transmission dynamics that are still not fully characterized. Future field applications will require transitioning from artificial injection models toward strategies that leverage robust oral infection alongside natural horizontal or vertical transmission. Furthermore, experience from genetically modified mosquito programs highlights the importance of regulatory frameworks, risk assessment, and public acceptance in determining feasibility(57). As such, further development of ISF-based strategies must integrate virological performance with ethical, regulatory, and societal considerations.

In summary, this research demonstrates that structural protein exchange improves oral infectivity and replication-dependent SIE in *Ae. aegypti*. While population-wide uniform suppression was not achieved, the association between high levels of chimera replication and a reduced number of arbovirus infections and reduced viral titres provides important mechanistic insight into the biological conditions under which ISF-mediated interference may occur. These findings extend the current understanding of mosquito vector, ISF and arbovirus interactions and define both the promise and the practical constraints of chimeric ISFs as tools for arbovirus control.

## Acknowledgments

The authors thank Corinne Geertsema for cell culture maintenance. We appreciate the assistance by dr. NCA de Ruijter and facilities provided by the Wageningen Light Microscopy Centre (WLMC). We thank Melissa Graham for her help with the optimisation during whole mosquito immunofluorescence staining (IFA) and Alexander W. E. Franz for his advice regarding the optimization of IFA for mosquito midguts.

## CRediT authorship contribution statement

**Wessel Willemsen:** Writing – original draft, Visualization, Validation, Methodology, Investigation, Funding acquisition, Formal analysis, Data curation, Conceptualization. **Alyssa J. Peterson:** Writing – original draft, Visualization, Validation, Methodology, Investigation, Formal analysis, Data curation, Conceptualization. **Marleen Henkens:** Investigation. **Hans Smid:** Writing –review & editing, Visualization. **Hayden J. Rohlf:** Investigation. **Tessa M. Visser:** Resources. **Constantianus J.M.Koenraadt:** Writing –review & editing, Resources. **Roy A. Hall:** Review, Supervision, Funding acquisition. **Jody Hobson-Peters:** Writing – review & editing, Supervision, Funding acquisition. **Monique van Oers:** Writing –review & editing. **Gorben P. Pijlman:** Writing – review & editing, Supervision, Funding acquisition, Conceptualization. **Jessica J. Harrison:** Supervision, Funding acquisition. **Leon E. Hugo:** Writing **-** review & editing Visualization, Supervision. **Jelke J. Fros:** Writing – review & editing, Supervision, Project administration, Funding acquisition, Data curation, Conceptualization.

## Declaration of generative AI and AI-assisted technologies in the writing process

During the preparation of this work the author(s) used ChatGPT, Gemini or Claude in order to improve language and readability. After using this tool/service, the author(s) reviewed and edited the content as needed and take(s) full responsibility for the content of the publication.

## Data availability

Small RNA sequencing libraries have been deposited in the NCBI Sequence Read Archive (SRA) under the associated BioProjectID:

## Supplemental Figures

**Figure S1. Validation of viral titers in infectious blood meals and blood-fed mosquitoes.** A, B) ZIKV titers in blood meals and whole mosquitoes collected immediately post-feeding. Panels show titers for the higher-dose blood meal (2 × 10^7^ TCID₅₀/mL) corresponding to the vector competence experiment in Figure 2 (A) and the lower-dose blood meal (2 × 10⁶ TCID₅₀/mL) corresponding to the experiment in Figure S2 (B). Data are shown across three independent replicates (R1–R3). C) Viral titers in blood meals and engorged mosquitoes corresponding to the midgut barrier analysis in Figure 4C. D) Viral titers in blood meals and engorged mosquitoes corresponding to the superinfection exclusion experiments shown in Figure 6. Data points represent individual blood meal batches or single mosquito samples, presented with their respective mean; error bars represent the standard error of the mean. The dotted line indicates the limit of detection (LoD).

**Figure S2. ZIKV infection and dissemination in mosquitoes injected with BinJV, BinJ-ZIKV following a lower-dose blood meal (2 × 10⁶ TCID₅₀/mL).** A) Experimental design. *Aedes aegypti* mosquitoes were first intrathoracically injected with either PBS (n=63), BinJV (n=39) or BinJ-ZIKV (n=38) (6.9 × 10^3^ TCID₅₀/mosquito). Four days later, mosquitoes received a blood meal containing ZIKV (2 × 10⁶ TCID₅₀/mL). At 10 days post ZIKV infection, bodies, legs and wings, and saliva were collected and infection prevalence was determined by infectivity assay after which positive samples were titrated by EPDA (B–D) Percentage of mosquitoes positive for ZIKV in bodies (B), legs and wings (C), and saliva (D) following prior injection with PBS, BinJV, or BinJ-ZIKV and subsequent infectious blood meal (2 × 10⁶ TCID₅₀/mL). (E–G) Individual mosquito ZIKV titres (log₁₀ TCID₅₀/mL) corresponding to B–D, with the limit of detection (LoD) indicated. Bars represent the average percentage of mosquitoes positive for ZIKV from three independent experiments. Asterisk indicates significant difference with the PBS injected group (GLM with replicate as blocking factor, p<0.05) The full statistical analyses are provided in the supplementary files (Table S1A,B). E-G) Data points represent viral titers obtained from a single mosquito sample. Individual samples from three biological replicates are presented together with their respective mean and standard error of the mean. The dotted line indicates the limit of detection (LoD).

---

**Table S1A.**
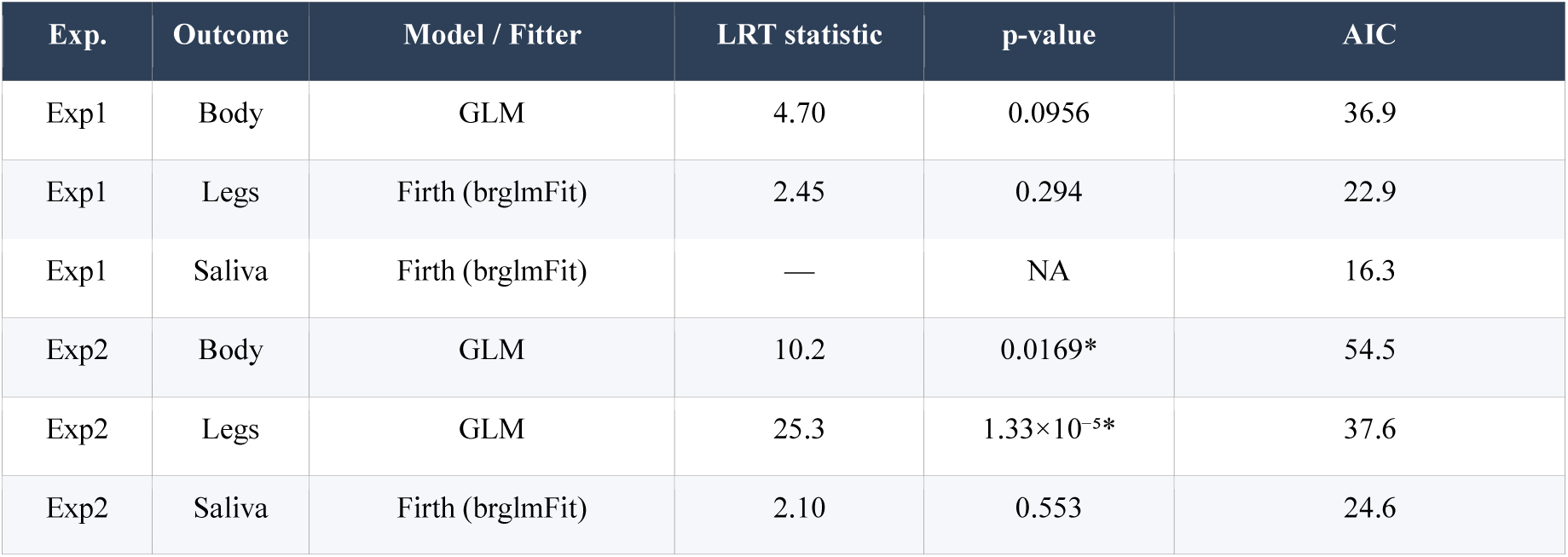
Overall likelihood ratio tests (LRT) for group effect per experiment and outcome.

**Table S1B.**
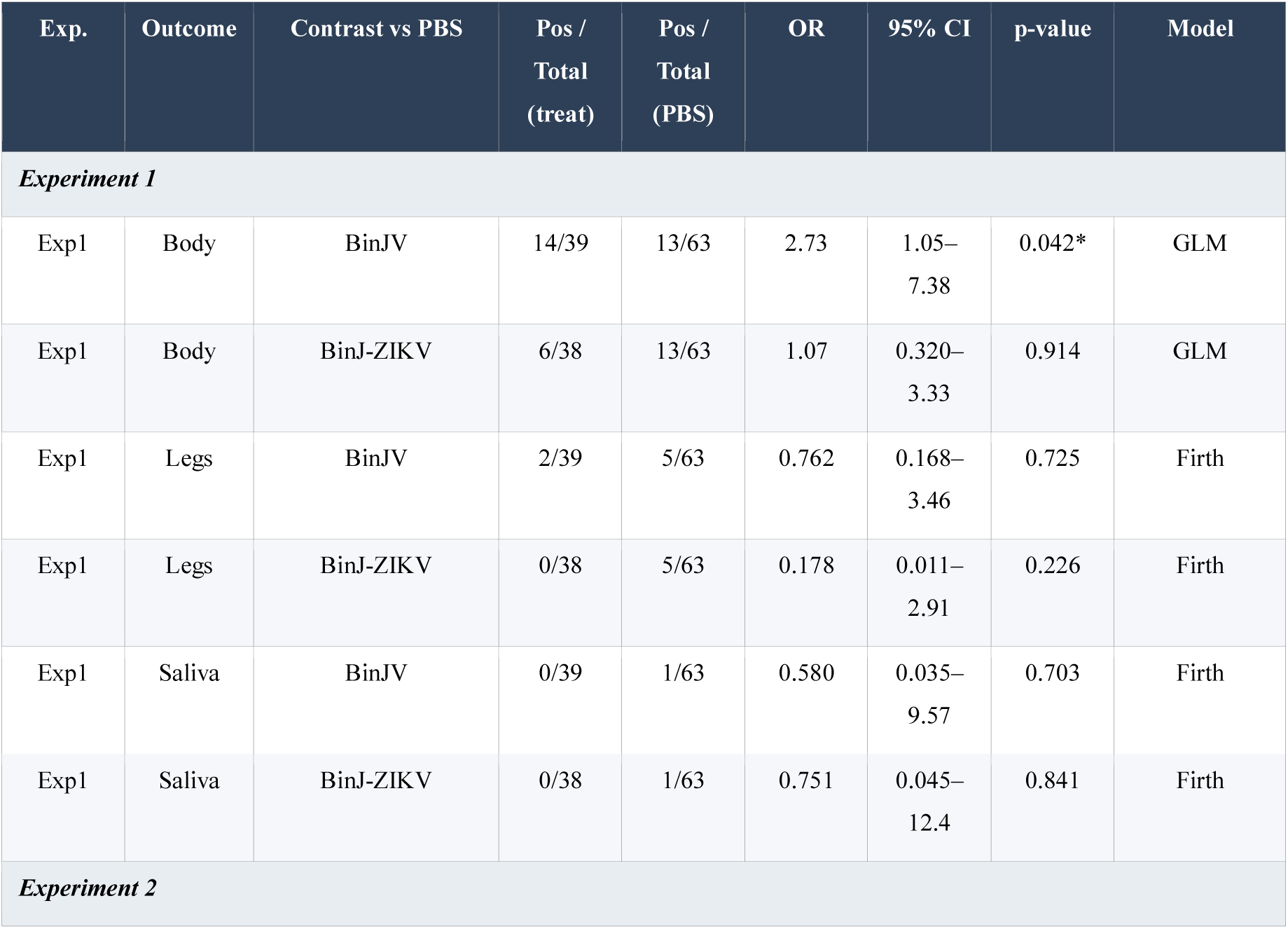
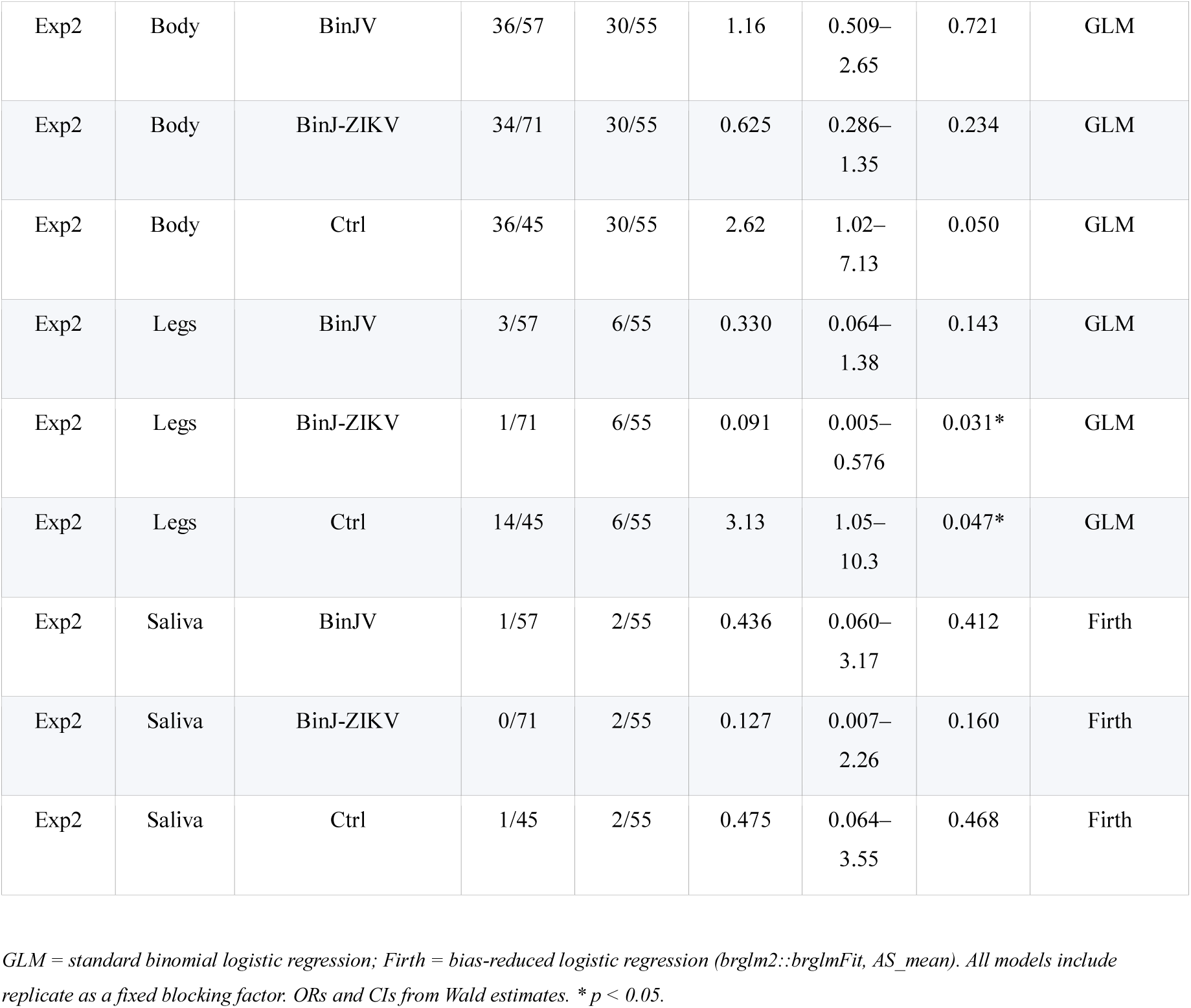
Pairwise contrasts vs PBS: odds ratios, 95% confidence intervals, and p-values.

**Table S2.**
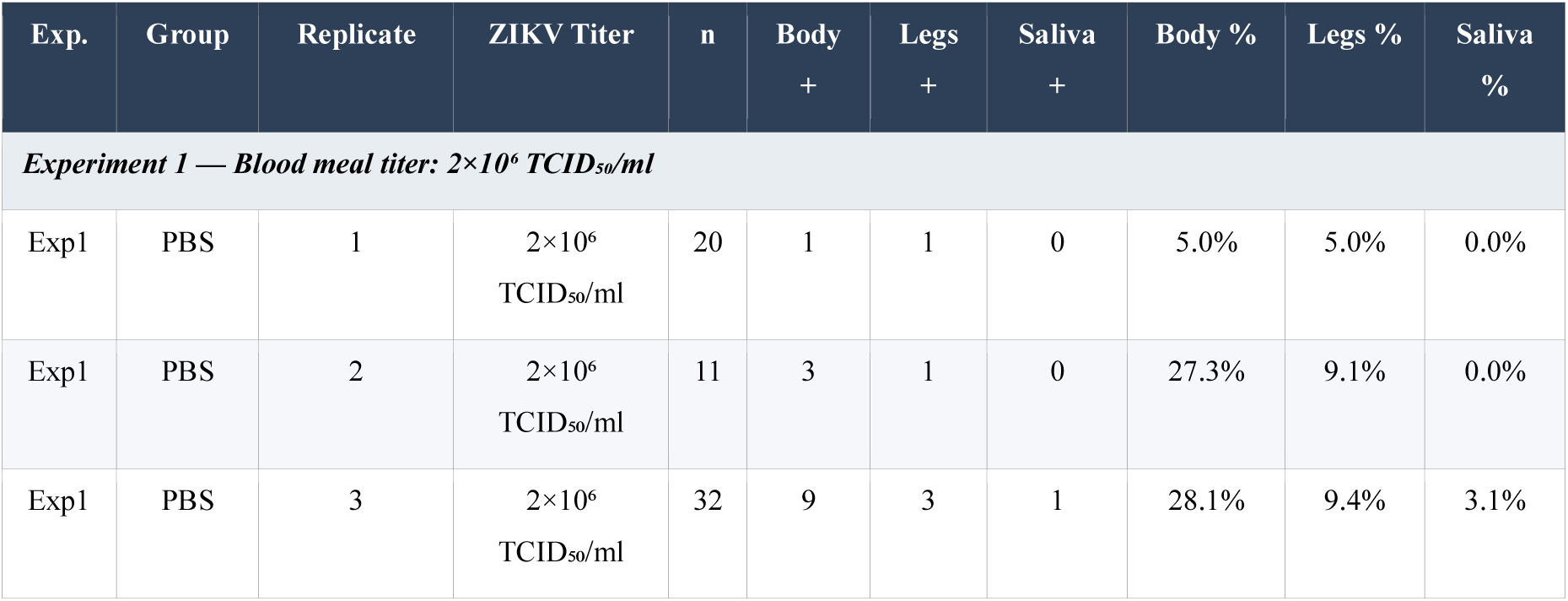
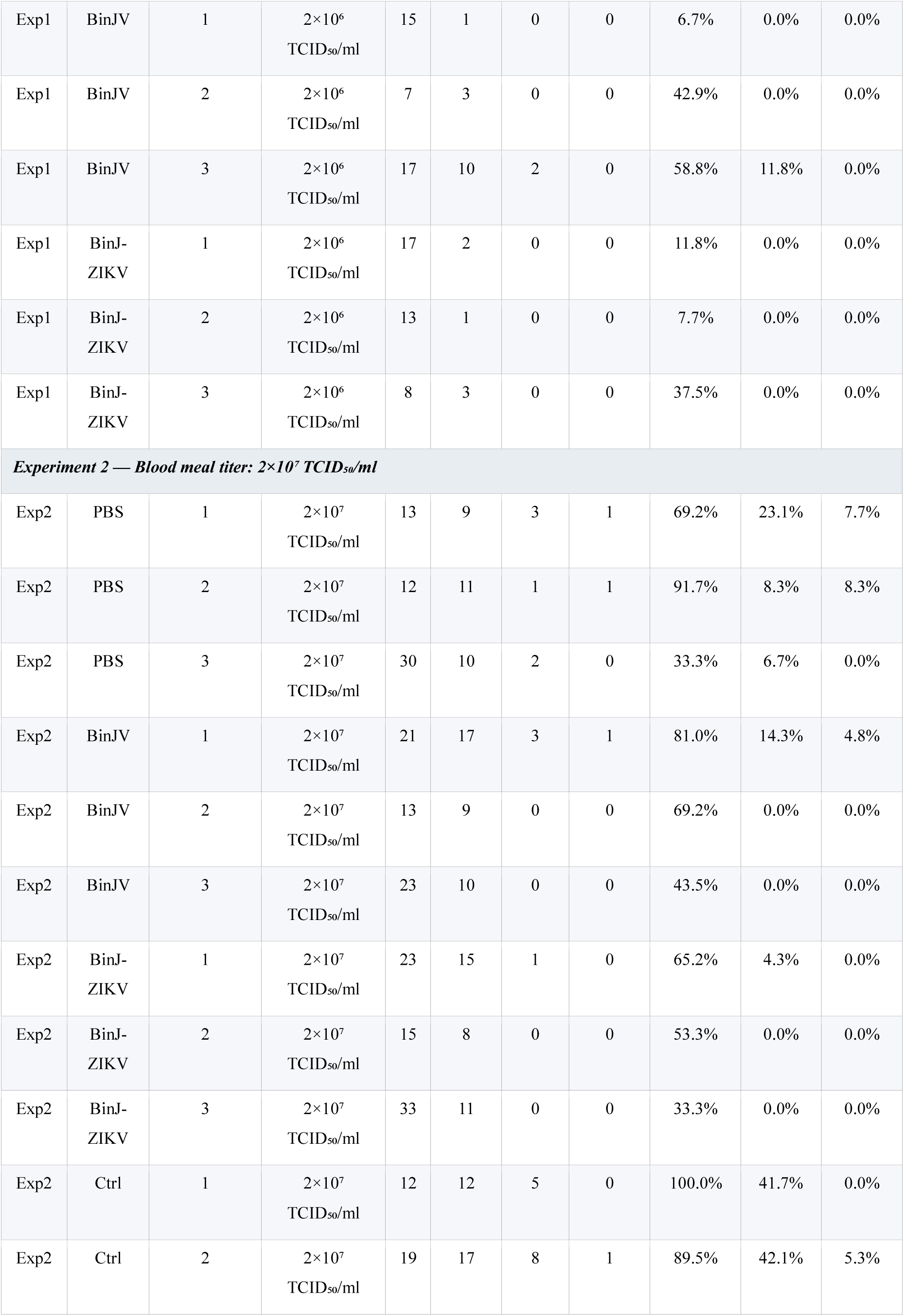
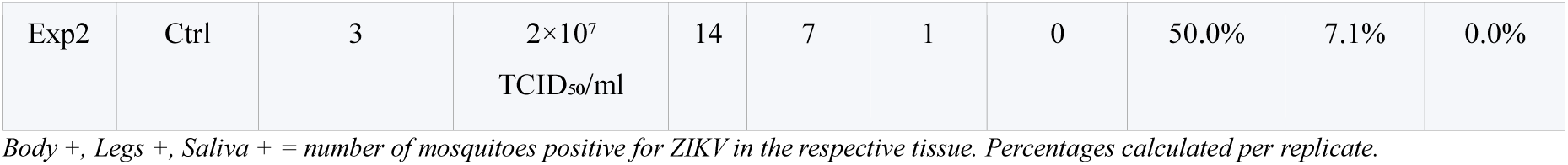
Raw mosquito counts and infection rates per replicate.

| Exp. | Group | Replicate | ZIKV Titer | n | Body<br>+ | Legs<br>+ | Saliva<br>+ | Body % | Legs % | Saliva<br>% |
| --- | --- | --- | --- | --- | --- | --- | --- | --- | --- | --- |
| *Experiment 1 — Blood meal titer: $2 \times 10^6$ TCID<sub>50</sub>/ml* |  |  |  |  |  |  |  |  |  |  |
| Exp1 | PBS | 1 | $2 \times 10^6$<br>TCID<sub>50</sub>/ml | 20 | 1 | 1 | 0 | 5.0% | 5.0% | 0.0% |
| Exp1 | PBS | 2 | $2 \times 10^6$<br>TCID<sub>50</sub>/ml | 11 | 3 | 1 | 0 | 27.3% | 9.1% | 0.0% |
| Exp1 | PBS | 3 | $2 \times 10^6$<br>TCID<sub>50</sub>/ml | 32 | 9 | 3 | 1 | 28.1% | 9.4% | 3.1% |
| Exp1 | BinJV | 1 | $2 \times 10^6$<br>TCID<sub>50</sub>/ml | 15 | 1 | 0 | 0 | 6.7% | 0.0% | 0.0% |
| Exp1 | BinJV | 2 | $2 \times 10^6$<br>TCID<sub>50</sub>/ml | 7 | 3 | 0 | 0 | 42.9% | 0.0% | 0.0% |
| Exp1 | BinJV | 3 | $2 \times 10^6$<br>TCID<sub>50</sub>/ml | 17 | 10 | 2 | 0 | 58.8% | 11.8% | 0.0% |
| Exp1 | BinJ-ZIKV | 1 | $2 \times 10^6$<br>TCID<sub>50</sub>/ml | 17 | 2 | 0 | 0 | 11.8% | 0.0% | 0.0% |
| Exp1 | BinJ-ZIKV | 2 | $2 \times 10^6$<br>TCID<sub>50</sub>/ml | 13 | 1 | 0 | 0 | 7.7% | 0.0% | 0.0% |
| Exp1 | BinJ-ZIKV | 3 | $2 \times 10^6$<br>TCID<sub>50</sub>/ml | 8 | 3 | 0 | 0 | 37.5% | 0.0% | 0.0% |
| **Experiment 2 — Blood meal titer: $2 \times 10^7$ TCID<sub>50</sub>/ml** |  |  |  |  |  |  |  |  |  |  |
| Exp2 | PBS | 1 | $2 \times 10^7$<br>TCID<sub>50</sub>/ml | 13 | 9 | 3 | 1 | 69.2% | 23.1% | 7.7% |
| Exp2 | PBS | 2 | $2 \times 10^7$<br>TCID<sub>50</sub>/ml | 12 | 11 | 1 | 1 | 91.7% | 8.3% | 8.3% |
| Exp2 | PBS | 3 | $2 \times 10^7$<br>TCID<sub>50</sub>/ml | 30 | 10 | 2 | 0 | 33.3% | 6.7% | 0.0% |
| Exp2 | BinJV | 1 | $2 \times 10^7$<br>TCID<sub>50</sub>/ml | 21 | 17 | 3 | 1 | 81.0% | 14.3% | 4.8% |
| Exp2 | BinJV | 2 | $2 \times 10^7$<br>TCID<sub>50</sub>/ml | 13 | 9 | 0 | 0 | 69.2% | 0.0% | 0.0% |
| Exp2 | BinJV | 3 | $2 \times 10^7$<br>TCID<sub>50</sub>/ml | 23 | 10 | 0 | 0 | 43.5% | 0.0% | 0.0% |
| Exp2 | BinJ-ZIKV | 1 | $2 \times 10^7$<br>TCID<sub>50</sub>/ml | 23 | 15 | 1 | 0 | 65.2% | 4.3% | 0.0% |
| Exp2 | BinJ-ZIKV | 2 | $2 \times 10^7$<br>TCID<sub>50</sub>/ml | 15 | 8 | 0 | 0 | 53.3% | 0.0% | 0.0% |
| Exp2 | BinJ-ZIKV | 3 | $2 \times 10^7$<br>TCID<sub>50</sub>/ml | 33 | 11 | 0 | 0 | 33.3% | 0.0% | 0.0% |
| Exp2 | Ctrl | 1 | $2 \times 10^7$<br>TCID<sub>50</sub>/ml | 12 | 12 | 5 | 0 | 100.0% | 41.7% | 0.0% |
| Exp2 | Ctrl | 2 | $2 \times 10^7$<br>TCID<sub>50</sub>/ml | 19 | 17 | 8 | 1 | 89.5% | 42.1% | 5.3% |
| Exp2 | Ctrl | 3 | $2 \times 10^7$<br>TCID<sub>50</sub>/ml | 14 | 7 | 1 | 0 | 50.0% | 7.1% | 0.0% |

*Body +, Legs +, Saliva + = number of mosquitoes positive for ZIKV in the respective tissue. Percentages calculated per replicate.*

